# Spread of the pandemic Zika virus lineage is associated with NS1 codon usage adaptation in humans

**DOI:** 10.1101/032839

**Authors:** Caio César de Melo Freire, Atila Iamarino, Daniel Ferreira de Lima Neto, Amadou Alpha Sall, Paolo Marinho de Andrade Zanotto

**Affiliations:** Laboratory of Molecular Evolution and Bioinformatics, Department of Microbiology, Biomedical Sciences Institute, University of Sao Paulo, Sao Paulo, Brazil.; Institut Pasteur de Dakar, Dakar, Senegal.

**Keywords:** Zika virus, emerging diseases, molecular evolution, codon usage adaptation, NS1

## Abstract

Zika virus (ZIKV) infections were more common in the zoonotic cycle until the end of the 20^th^ century with few human cases in Africa and Southeastern Asia. Recently, the Asian lineage of ZIKV is spreading along human-to-human chains of transmission in the Pacific Islands and in South America. To better understand its recent urban expansion, we compared genetic differences among the lineages. Herein we show that the recent Asian lineage spread is associated with significant NS1 codon usage adaptation to human housekeeping genes, which could facilitate viral replication and increase viral titers. These findings were supported by a significant correlation with growth in Malthusian fitness. Furthermore, we predicted several epitopes in the NS1 protein that are shared between ZIKV and Dengue. Our results imply in a significant dependence of the recent human ZIKV spread on NS1 translational selection.

## INTRODUCTION

Changes in nucleotide composition have long been noticed as an important evolutionary mechanism and a telltale of viral adaptation to host (Pepin et al., 2010; Plotkin and Kudla, 2011; Longdon et al., 2014). Codon usage adaptation after a host shift event could be required to fine-tune the interactions between a virus and a new host (Longdon et al., 2014; Bahir et al., 2009). Zika virus (ZIKV) was known as a zoonotic pathogen with sporadic human infections in Africa and latter in Southeastern Asia until the end of the last century (Hayes, 2009). In Africa, it remains in a sylvatic cycle involving mainly monkeys and several *Aedes* mosquitoes (Faye et al., 2014). While its Asian lineage is spreading along long chains human-to-human transmission in the Pacific Islands and in South America, vectored mainly by *Aedes aegypti* (Musso et al., 2015). Crucially, the ZIKV pandemic potential is maximized by being also vectored by *A. albopictus* (Grard et al., 2014), a mosquito that explores higher latitudes and transmitted Chikungunya virus in USA and Europe recently (Kuehn, 2014; Grandadam et al., 2011; Delisle et al., 2015). Additionally, sexual intercourse and perinatal infection may be alternative routes of transmission (Besnard et al., 2014; Foy et al., 2011).

The Asian lineage first caused an outbreak of febrile disease in Yap Island, Federated States of Micronesia, in 2007 (Duffy et al., 2009; Hayes, 2009). In 2013 and 2014, it emerged again and caused a large epidemic in French Polynesia (Cao-Lormeau et al., 2014), spreading to Oceania and arriving in America at Easter Island by 2014 (Musso et al., 2015). Recently, in early 2015, it was reported in several Brazilian provinces (Zanluca et al., 2015; Campos et al., 2015), mainly in the Northeastern region. The intense tourism in this regions promotes a massive traffic of people between Brazil and Europe and could help spread ZIKV further, such as when a traveler returned with Zika fever (ZF) from Bahia state in the Northeastern of Brazil to Italy (Zammarchi et al., 2015). ZF symptoms include lasting arthralgia, headaches and mild fever (Zanluca et al., 2015; Campos et al., 2015). The recent outbreaks of ZIKV infections were also associated with a 20-fold increase in Guillain-Barre syndrome cases in French Polynesia (Musso et al., 2014). The increasing in Guillain-Barre cases was also observed in Bahia state, where ZIKV transmission is concomitant with Dengue (DENV) and Chikungunya viruses (CHIV) and ZF incidence reached 275 cases per 100,000 inhabitants until August 2015 (SESAB, 2015). Worryingly, ZIKV was recently associated to the abrupt increase of newborns with microcephaly in Brazil (Ministério da Saúde, 2015).

## RESULTS

**Codon preferences of ZIKV lineages are distinct** Because codon preferences can strongly affect gene expression (Plotkin and Kudla, 2011), we estimated the relative synonymous codon usage (RSCU) values (Sharp et al., 1986) for each ZIKV gene sequence (Figure 1A). By means of a principal component analysis (PCA) for RSCU values, we found distinct codon preferences in the African and Asian lineages for the entire polyprotein (Figure 1A) and for each viral gene (Figure S1). The extent of the codon bias was inferred by plotting the effective number of codons versus the proportion of GC-content in the third position for each codon (Wright, 1990). As a consequence, we found significant codon usage bias under purifying selective pressure (Wright, 1990), constraining the codon usage in ZIKV (Figure S2A and Figure S3), as found for other arboviruses (Jenkins and Holmes, 2003). The strong purifying selection, which we found at several codon sites (Table S1), was also observed for mosquito transmitted viruses that cause acute infections, mainly alternating between vectors and vertebrate hosts (Hanada et al., 2004). As expected, high amino acid conservation was observed; *e.g*. 91.8% of the 353 residues of the 17 NS1 proteins analyzed were identical, which is indicative of purifying selection.

**Figure 1.**
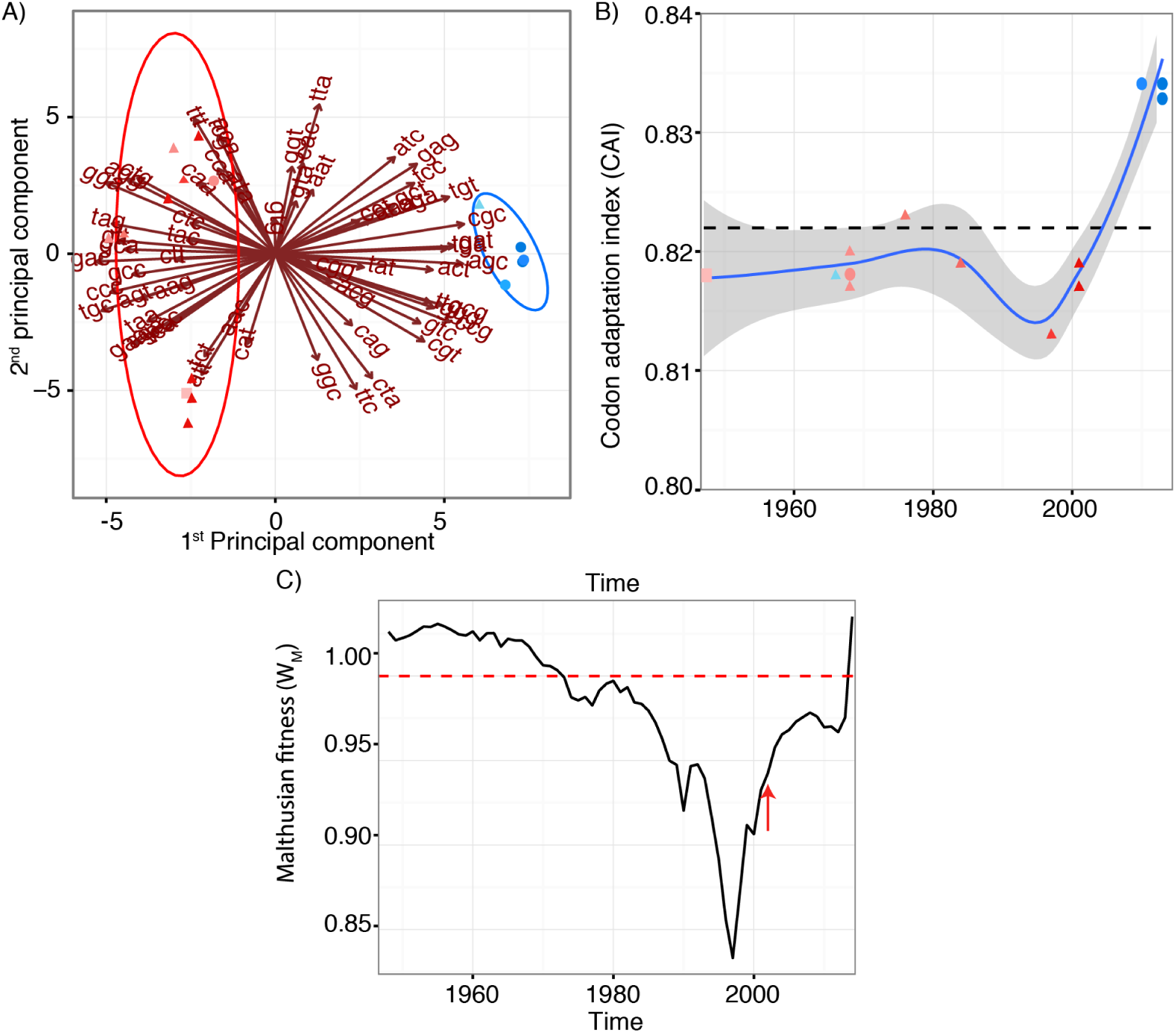
Zika Virus (ZIKV) codon adaptation and fitness according to lineage. A) RSCU analysis for the polyprotein coding region shows that the principal component (PCA) agrees with the phylogenetic distinctions between the two ZIKV lineages. The African (red) and Asian (blue) lineages were color-coded according to the isolation dates, lighter colors represent older isolation dates. Shapes represent the isolation host: mosquitoes (triangle), monkey (square) or humans (circles). B) NS1 gene Codon Adaptation Index (CAI) to the human housekeeping genes for the African (red) and Asian (blue) lineages according to the isolation dates. C) Malthusian fitness (*W_M_*) estimated for ZIKV since 1947, representing decrease (*W_M_* < 1), constant population size (*W_M_* = 1), and net growth (*W_M_* > 1). The red arrow references the end of African lineage sampling. The Spearman correlation coefficients (*ρ*) between the interpolated CAI values in the Figure 1B and the estimated *W_M_* in Figure 1C were calculated for three time periods: (i) 1948–1970, (*ii*) 1971–1992 and (*iii*) 1992–2014. For the former period, we observed a significant negative correlation (*ρ* = −0.59 and *ρ*-value = 0.004); in the second the correlation was significant and positive (*ρ* = 0.46 and p-value = 0.04) and in the most recent period we found a significant strong positive correlation (*ρ* = 0.90 and p-value = 2.70*E –* 6).

**Evidences of translational selection to human codon usage in NS1 gene of ZIKV epidemic lineages**. Given the significantly distinct codon preferences between the African and Asian ZIKV lineages shown in Figure 1A, and because only the Asian lineage is associated to massive human outbreaks recently (Musso et al., 2015), we further compared virus codon adaptation to host and vector, human housekeeping and *Aedes aegypti* genes. We found that recent Asian epidemic lineages had stronger codon bias on NS1 and NS4A genes, and were also more adapted to humans. This was suggested by measurements of the codon adaptation index (CAI) (Sharp and Li, 1987) for each ZIKV gene for both lineages, unveiling potential viral adaptation to cellular translation machinery of man and mosquitoes. We show in Figure S2B that all ZIKV strains were significantly adapted to humans (CAI values above threshold for the entire polyprotein) while less adapted to *Aedes aegypti* mosquitoes. Moreover, when CAI values were calculated separately for most genes there was little differences between lineages (Figure S4), as was expected for most of the viral genes (Bahir et al., 2009). Nevertheless, codon adaptation for humans in the NS1 coding region from the recent Asian lineage showed a clear increase in CAI values near the present (Figure 1B), coinciding with its spread to Pacific and America. The strong bias in codon usage observed for NS1 from epidemic strains provided additional evidence of translational selection acting on this gene (solid blue circles in Figure S3D).

**High CAI values for the Asian lineage were correlated with the recent ZIKV growth**. This relevant finding was supported by both, (*i*) the concurrent Malthusian fitness (*W_M_*) values above one (Day and Otto, 2001) and, (*ii*) the significant strong positive correlation (*ρ* = 0.90 and p-value = 2.70*E –* 6) between *W_M_* and the interpolated values for CAI in the period from 1990 to 2014 (Figure 1B and 1C). Viral codon usage optimization is critical for fine-tuning the interaction with a given host (Longdon et al., 2014), and the most affected genes are usually those highly expressed (Bahir et al., 2009). Therefore, the high NS1 CAI values that we observed for recent Asian ZIKV were considered as a strong indication of adaptive change, which could be associated to improvement in translational efficiency in humans and increased viremia in patients, as observed for Lassa virus (Andersen et al., 2015). Moreover, the NS1 protein is secreted at high levels by infected cells as hexamers that are implicated in immune evasion strategies (Muller and Young, 2013). We obtained similar results on translational selection on NS4A (Figure S3H and S4H). This may be relevant, since NS4A and NS1 appear to play a role in viral replication (Lindenbach and Rice, 1999), while NS4A may enhance viral survival by preventing cell death by the up-regulation of cell autophagy (McLean et al., 2011).

**ZIKV and DENV could share B-cell epitopes**. Another relevant function of the NS1 protein is in assisting in flavivirus immune evasion (Muller and Young, 2013). NS1-specific antibodies are usually found during secondary infections and there is NS1 cross-reactivity between ZIKV and DENV (Lanciotti et al., 2008; Valdés et al., 2000; Muller and Young, 2013), which could impact on pathogenesis. Because *in silico* epitope prediction have been used extensively to develop peptide-based vaccines and investigate immune responses (He and Zhu, 2015), we inferred the structural similarity between the NS1 of DENV and ZIKV by homology modeling (Figure S5A). We found nine linear and five discontinuous epitopes shared in equivalent positions, despite low sequence identity among them (Table S2). Nevertheless, linear epitopes also shared physicochemical properties (Figure S5B). We further calculated the root mean square deviation (RMSD) and performed the global distance test (GDT) for the shared conformational epitopes and found that they were structurally similar, which reinforce the notion that these epitopes may be shared in these phylogenetically closely related viruses (Kuno et al., 1998). These findings could explain the observation of aggravated health conditions in co-infections or secondary infections by ZIKV on DENV pre-exposed people (Roth et al., 2014).

## DISCUSSION

The differences between the African and Asian lineages could explain the emergence of ZIKV in humans and raises concerns about the consequences of the adaptive genetic changes observed in NS1 (Figure 1B) and the recent increase in viral fitness (Figure 1C) (Pepin et al., 2010; Longdon et al., 2014). Moreover, the limited number of human ZIKV cases in Africa could be associated to low viremia in humans, which was demonstrated by a health-officer volunteer experimentally infected with a virus from the African lineage that failed to infect A. *aegypti* mosquitoes (Bearcroft, 1956). Together, our results suggest that fitness gain is associated with improvement of the NS1 translation in humans by synonymous mutations. Synonymous mutations are a common source of variation, given the constrained non-synonymous substitutions rate imposed to RNA viruses that have to negotiate successful infections, alternating between humans and mosquitoes (Hanada et al., 2004). It remains to be evaluated how the NS1 structural and immunological similarities associate to the aggravated symptoms observed when ZIKV and DENV co-circulate (Roth et al., 2014). For this reason, our findings may also be of considerable relevance for the ongoing development of DENV vaccines (McArthur et al., 2013).

## METHODS

**Sequence datasets**. We investigated all 17 available complete genome sequences of ZIKV from GenBank that had information of year and country of isolation (alignment available in https://github.com/CaioFreire/CUB). First, we aligned the coding sequences with MACSE program v0.9 (Ranwez et al., 2011) and curated it with SeaView v4.40 (Gouy et al., 2010). During phylodynamic analyses, we employed the most comprehensive dataset for 51 NS5 gene sequences (also available in https://github.com/CaioFreire/CUB) sampled from 1947 to 2014, ranging from 14 countries in Africa, Asia, Pacific Islands and America. Since we previously found evidences of recombination in ZIKV from Africa (Faye et al., 2014) and these events could cause potential errors in phylogenetic inferences (Posada and Crandall, 2002), we screened for recombination in NS5 sequences with the RDP program v4.36 (Martin et al., 2010). Identified recombinants were removed of phylogenetic-based analysis. Codon preferences analyses. We employed the relative synonymous codon usage method (Sharp et al., 1986) with the R-package SeqinR v3.13 (Charif and Lobry, 2007) to estimate the codon preferences for each polyprotein gene sequence. In addition, we employed a principal component analysis (PCA) to assess patterns among RSCU values among viral lineages (Su et al., 2009). We identified the most informative codons, which were informative to discriminate among Asian and African lineages, with a biplot graph for the PCA values with the R-package ggbiplot v0.55 (Vu, 2011), using a group probability of 0.95. The different codon preferences between ZIKV lineages were independently confirmed by high support values (> 80%) obtained from hierarchical clustering analysis, using the R-package Pvclust v1.32 (Suzuki and Shimodaira, 2006).

**Codon usage biases**. We calculated the effective number of codons (ENC) with Emboss v6.60 (Rice et al., 2000) and the proportion of guanine-cytosine content in the third base of the codons (GC3), using Seqin{R} program to evaluate the codon usage bias (CUB). The theoretical curve of ENC x GC3 on the genetic drift was estimated with a Perl script to calculate expected ENC and GC3 values (available in https://github.com/CaioFreire/CUB), according to (Wright, 1990).

**Codon adaptation of ZIKV genes to humans and *Aedes aegypti* mosquitoes**. In our analysis, CAI is a measure of synonymous codon usage bias based on the codon preference of a viral strain and a codon usage table for a given host (Sharp et al., 1986). To investigate if the codon usage of ZIKV lineages was similar to the hosts in urban settings, regarding humans and *Aedes aegypti*, we calculated the codon adaptation indices (CAI) for each gene from each ZIKV lineage. Since the most pronounced biases are in highly expressed genes (Sharp and Li, 1987; Bahir et al., 2009), we used Emboss to calculate a codon usage table for humans (available in https://github.com/CaioFreire/CUB) based on 3803 genes identified as housekeeping (Suzuki and Shimodaira, 2006). Moreover, we calculated CAI for A. *aegypti* using the table available in Codon Usage Database (Nakamura et al., 2000). Importantly, the CAI values obtained with our table based on housekeeping genes were very similar to those found with the table from Codon Usage Database with generic human genes. The CAI values for each sequence from ZIKV genes were calculated with CAIcal program (Puigbò et al., 2008). We assessed the confidence of CAI estimates by the calculation of expected CAI values for 500 random sequences with similar GC-content and codon composition for each gene.

**Selection analyses**. We investigated the selection regimens acting on the polyprotein codon sites, calculating the difference (*ω*) between the estimates of non-synonymous *(dN)* and synonymous (*dS*) substitution rates per codon site. The *ω* values were estimated with single likelihood ancestor counting (SLAC) method with HyPhy program v2.11 (Pond et al., 2005), assuming a significance level (α) of 0.05. We employed a maximum likelihood (ML) phylogenetic tree, inferred with GARLI v2.01 (Zwickl, 2006), on NS5 gene alignment without recombinant sequences and the polyprotein gene alignment for the taxa without recombination in the NS5 gene, as input to SLAC. Codon sites under purifying selection were revealed by *ω* < 0, and the opposite is indicative of diversifying selection.

**Phylodynamic analyses**. Using dates of isolation, we were able to estimate a time-scaled Maximum Clade Credibility (MCC) tree for ZIKV NS5 sequences (alignment available in https://github.com/CaioFreire/CUB). We used BEAST v1.82 (Drummond et al., 2012), with the evolutionary rate prior *(μ*) of 1*x*10 – 3 found previously (Faye et al., 2014). Since purifying selection could underestimate the time to the most recent common ancestor (TMRCA) (Wertheim and Kosakovsky Pond, 2011), we used a substitution model for protein-coding sequences (SRD06) (Shapiro et al., 2006). To infer the demographic history of ZIKV, we employed the Bayesian skyride method (Minin et al., 2008) to estimate the temporal dynamics of effective population size (*Ne.g*) of ZIKV, which approximates the number of infections in time. To reveal the dynamics of viral population size growth, we calculated the Malthusian fitness *(W_M_*), which was approximated by the ratio of the population size in sequential time points (*W_M_* = *Ne.g_t_/Ne.g_t_* – 1) (Day and Otto, 2001). Moreover, we investigated the correlations between interpolated CAI values for NS1 and *W_M_*, using the Spearman rank correlation tests in three time intervals: (*i*) 1947–1969, (*ii*) 1970–1990, and (*iii*) 1991–2014.

**Homology modeling and Linear / Discontinuous Epitope Prediction**. These analyses were based on references sequences, available in GenBank, of NS1 protein from Dengue subtypes 1 to 4 (GenBank accession numbers: AGN94879, AGN94890, ABV03585 and AFX65881) and from ZIKV strains from Senegal and French (GenBank accession numbers: AEN75266.1 and AHZ13508). We aligned the sequences with AliView (Larsson, 2014) and MUSCLE (Edgar, 2004). Sequences with less than 95% identity were selected for each subtype and modeled using YASARA (Krieger and Vriend, 2015) in a BioLinux 8 (Afgan, 2012) with 20 PSI-Blast iterations (e-value = 0.7), considering 6 oligomerization states. 20 templates were downloaded from the Protein Data Bank (PDB - http://www.rcsb.org/pdb/) with 5 sequence alignments per template. Modeling was set to low speed with 10 terminal extensions, sampling 50 terminal loops. We checked the produced structures for consistency at the PDBSum server (de Beer et al., 2014) with the Generate option (available at https://www.ebi.ac.uk/thornton-srv/databases/cgi-bin/pdbsum/) and PROCHECK (Laskowski et al., 1993) stereo-chemical analyses. We calculated relative accessible surface area using the modeled structures with the server GETAREA (Fraczkiewicz and Braun, 1998) (available at http://curie.utmb.edu/getarea.html). Linear and discontinuous B Cell epitopes were predicted using the Immune Epitope Database (http://tools.immuneepitope.org/). We used the module YASARA view to map on the modeled structures the epitopes found by the IEDB server (Haste Andersen et al., 2006). Linear epitopes were predicted using the Bepipred Linear Epitope Prediction (Larsen et al., 2006). Structural alignments were made to evaluate RMSD and GDT scores between the models for the epitope regions. All results are available in https://github.com/CaioFreire/CUB.

## Author contributions

CCMF, AI, DFLN, AAS, and PMAZ designed the experiments and wrote the paper. CCMF and DFLN conducted the experiments. CCMF, AI and DFLN prepared the figures.

## Acknowledgments

We thank the Fundação de Amparo à Pesquisa do Estado de São Paulo (FAPESP) for the funding (project #2014/17766–9). CCMF and AI also thank FAPESP for scholarships (#2012/04818–5 and #2014/06090–4). PMAZ holds a CNPq scholarship.

